# Fluximplied: A novel approach integrates rate limiting steps and differential expression for pathway analysis

**DOI:** 10.1101/2022.03.15.484417

**Authors:** Mike Sportiello, Adam Geber, Rohith Palli, A Karim Embong, Nathan G. Laniewski, Emma C. Reilly, Kris Lambert Emo, David J. Topham

## Abstract

**Background:** Many tools exist to perform a rigorous pathway analysis, though traditional gene set enrichment analysis remains among the most common. While useful for many applications, one common situation where it is less so is the metabolic profiling of bulk omics datasets. Rate limiting steps in more linear pathways are the main determinant of flux through these pathways, but differential expression of the enzymes that catalyze these steps is usually not differentially weighted in pathway analysis. Fluximplied was built to perform pathway analysis with rate limiting steps in mind to assess the implied flux through a number of well validated metabolic pathways.

**Results:** A database of rate limiting steps and their associated pathway was constructed. Using publicly available human RNA sequencing data from liver, putamen, and adipose tissue, fluximplied generally corroborated the pathway analysis. When comparing two CD8 T cell subsets from mouse lung, our algorithm confirmed previous findings that were not found previously.

**Conclusion:** Fluximplied is an accessible tool for pathway analysis which is intended to assist the user with hypothesis generation. Unlike traditional approaches to pathway analysis, it specifically queries a database of rate limiting steps in order to infer flux through canonical metabolic pathways.

## Background

Pathway analysis is a method that more deeply characterizes biology at the systems level. Pathway analysis generally presumes that the regulation of a single element (e.g. transcripts in RNA sequencing (RNAseq)) is less important than that of the pathway of interest as a whole. Traditional gene set enrichment analysis (GSEA) uses a list of differentially expressed transcripts (upregulated or downregulated) to assess if transcripts of a particular pathway are enriched in those lists. For example, if all elements of a pathway for *T cell killing* were upregulated but one, most would take this to be a biologically meaningful finding, even though one element is not. Pathway analysis by gene set enrichment analysis (GSEA), where enrichment of a predefined functional set of genes (*T cell killing* genes, for example) within a set of user-supplied genes (such as a list of differentially expressed genes from an RNAseq experiment), is assessed using the hypergeometric test.

More advanced methods make use of the particular amount of dysregulation of each gene as well as the underlying known regulatory topology of the network^1–9^. One popular and easy-to-use functional enrichment approach, EnrichR, uses the Fisher exact test to quantitatively average differences across a pathway. The recent Boolean Omics Network Invariant-Time Analysis (BONITA) uses Boolean approximation of flow through a network to give extra weight to elements with the highest impact ^10^. While these approaches are suitable for signaling cascades, no methods utilize knowledge of chemical equilibria or rate constants to improve pathway analysis in metabolic pathways.

Here we present fluximplied: a novel, free and open source software for pathway analysis and hypothesis generation. Fluximplied uses a manually curated database of rate limiting steps to predict increases or decreases in flux in a known metabolic pathway using transcriptomic data provided by the user.

Fluximplied can be used to generate hypotheses in the context of metabolic pathway analysis after creating a list of up or down-regulated genes derived from data generated using RNAseq, ATACseq, and other omics technologies. In GSEA, a list of upregulated genes is generated by using a Log_2_(Fold Change) (LFC) cutoff value to ensure changes are biologically meaningful and an adjusted p value threshold. In glycolysis for example, this type of analysis treats the upregulation of enolase—an enzyme that has little to do with either the regulation or rate of the pathway—the same as phosphofructokinase, which determines the rate and therefore the flux of the pathway as a whole. All else held equal, if the concentration of the RLS increases, the flux through the pathway will increase ^11^. Not knowing if all else is held equal, an increase in RLS *implies* increased flux through the pathway. It is important to note that a singular, pure RLS is rare: more often the rate is impacted to some degree by more than one enzyme in a pathway. However, more complicated methods like metabolic control analysis are quantitatively difficult, tissue dependent, and difficult to integrate with other methods like RNAseq. Usually one of only a small number of enzymes impact the rate to a meaningful degree, so the model of a singular RLS is still a useful one in analyses and hypothesis generation^12^.

Thus, we hypothesized the case where only a small number of genes could be upregulated and not yield a p value below the α threshold of 0.05, but where the upregulated genes encode RLS enzymes and were therefore the strongest determinant of flux. In **Figure 1**, three scenarios are presented. In all three scenarios, 100 genes are present, 10 of which are members of gene set *Green.* All three scenarios have 10 upregulated genes, 10 downregulated genes, and 80 of which are not differentially expressed. Since there are 10 upregulated genes, or 10% of the total genes, we would expect a non-enriched gene set of equivalent size to have 10% of its genes present (i.e. one gene). In Scenario A, we see there are 8 genes present: *Green* is enriched in our upregulated genes. In the second Scenario B, we have the same number of genes, but the RLS is not differentially expressed. In this example, *Green* is still enriched. In the final Scenario C, *Green* is not enriched, but there is likely a meaningful increase in flux through this pathway. It is this final scenario where GSEA and other traditional pathway analysis approaches are prone to false negatives.

**Figure 1:**
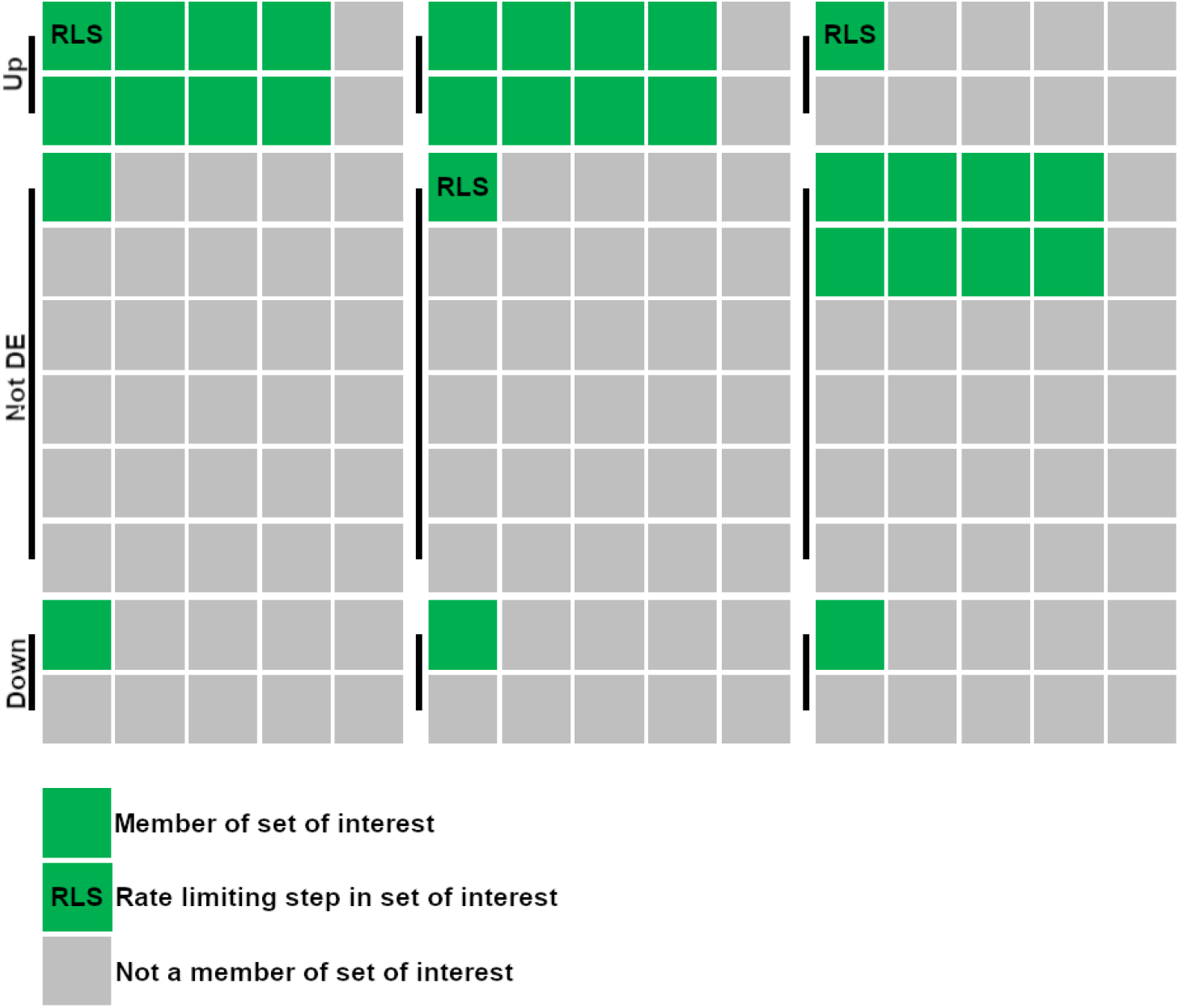
Situations of utility for fluximplied. Three scenarios are portrayed illustrating the situations where fluximplied would be more powerful than simple gene set enrichment analysis.

Furthermore, traditional enrichment analysis is agnostic of the LFC between conditions (unless an LFC cutoff is used) and thus a gene that has an LFC of 0.5 would be treated the same as a gene that has an LFC of 5, even though one has a fold change of 1.4 and the other 32. While this method is often useful and yields biologically meaningful results, it does not allow the user to more fully interrogate their data.

To that end, we developed fluximplied: the accessible, free and open source software to assist the user in generating hypotheses in metabolic pathway analysis and assess implied flux, a method typically requiring a time-consuming, large-scale flux balance analysis. Because the software was designed with bioinformatic novices in mind, a web-hosted interactive graphical user interface (GUI) was designed, although the tool remains powerful for all users in its capacity to integrate into computational pipelines. Fluximplied returns a differential expression analysis table of tested RLSs (with assigned adjusted p values and the corresponding pathways) and plots this table for visualization, all of which is downloadable. If the user is not able to access the differential expression analyses to feed into the software, they may supply a set of genes to fluximplied. In that scenario, since there is data like P values or LFCs available for generating a hypothesis if the user only supplies a list of genes, fluximplied matches any RLS available in such a list to pathways that may be impacted, returning a list of pathways the user may wish to further interrogate.

## Implementation

Fluximplied is written in R to be easily integrated into standard of the field omics analysis pipelines, though it calls upon a public rate limiting step database, which is maintained as a CSV. GitHub hosts the RLS database, and users can interface with the public repository to upload their own curated RLS databases for public use.

It can be used as a web-based application or integrated into standard RNAseq analysis pipelines through an R package. All of the authors participated in testing of the web-based app in different browsers and operating systems. Instructions for install and usage are kept most up to date at https://github.com/sportiellomike/fluximplied, though current instructions for install are below.

### R package

#### Install

The R package can be access through the R command line:

~~~
install.packages(’devtools’)
library(devtools)
install_github(’sportiellomike/fluximplied’,build_vignettes=T)
library(fluximplied)
~~~

#### Inputs

Once installed and loaded, the fluximplied’s main function fluximplied() takes six main arguments:

- Inputdat: takes two forms: a character vector of genes, or a data frame with gene names as row names as well as a column titled ‘log2FoldChange’ and a column of p values that the user can specify by name with the argument “padjcolname =” (**Table 1**). The format is specified under ‘inputformat’.
- species: takes ‘mmu’ or ‘hsa’ currently, referring to mouse or human respectively
- geneformat: takes ‘Symbol’ or ‘ENTREZ’ currently
- inputformat: the user must specify either ‘vector’ to refer to the character vector of genes supplied, or ‘dataframe’, which must include genes as row names as well as a column ‘log2FoldChange’ and one for adjusted P values
- padjcolname: the user must supply the name of the column where the adjusted P values are so that adjusted adjusted P values (P_adjadj_) can be calculated.
- pcutoff: the significance threshold for P_adjadj_, over which results will not be included in outputs. Defaults to 0.05.

**Table 1:** Data frame format for inputdat Correctly formatted inputdat of format ‘data frame’ that fluximplied will parse and analyze

#### Outputs

The outputs of fluximplied supplied with a data frame of information include

- A significance table which includes the significant metabolic pathways and their KEGG Pathway ID and P_adjadj_. These are appended to the supplied data frame, which would include at least the gene name, Padj, and log_2_(fold change) as well (**Table 2**).
- A plot of each significant pathway’s RLS’ Log_2_(Fold Change) colored by P_adjadj_ (**Figure 2**)
- A text explanation to aid in interpretation of these results.
- A character vector of the RLS genes included in your supplied data called myRLSgenes

**Table 2:** Example significance table output Fluximplied() will return a significance table which includes the supplied columns for gene names, P values, log_2_(fold change), as well as the calculated P_adjadj_, metabolic reaction pathway, and KEGG Pathway ID

**Figure 2:**
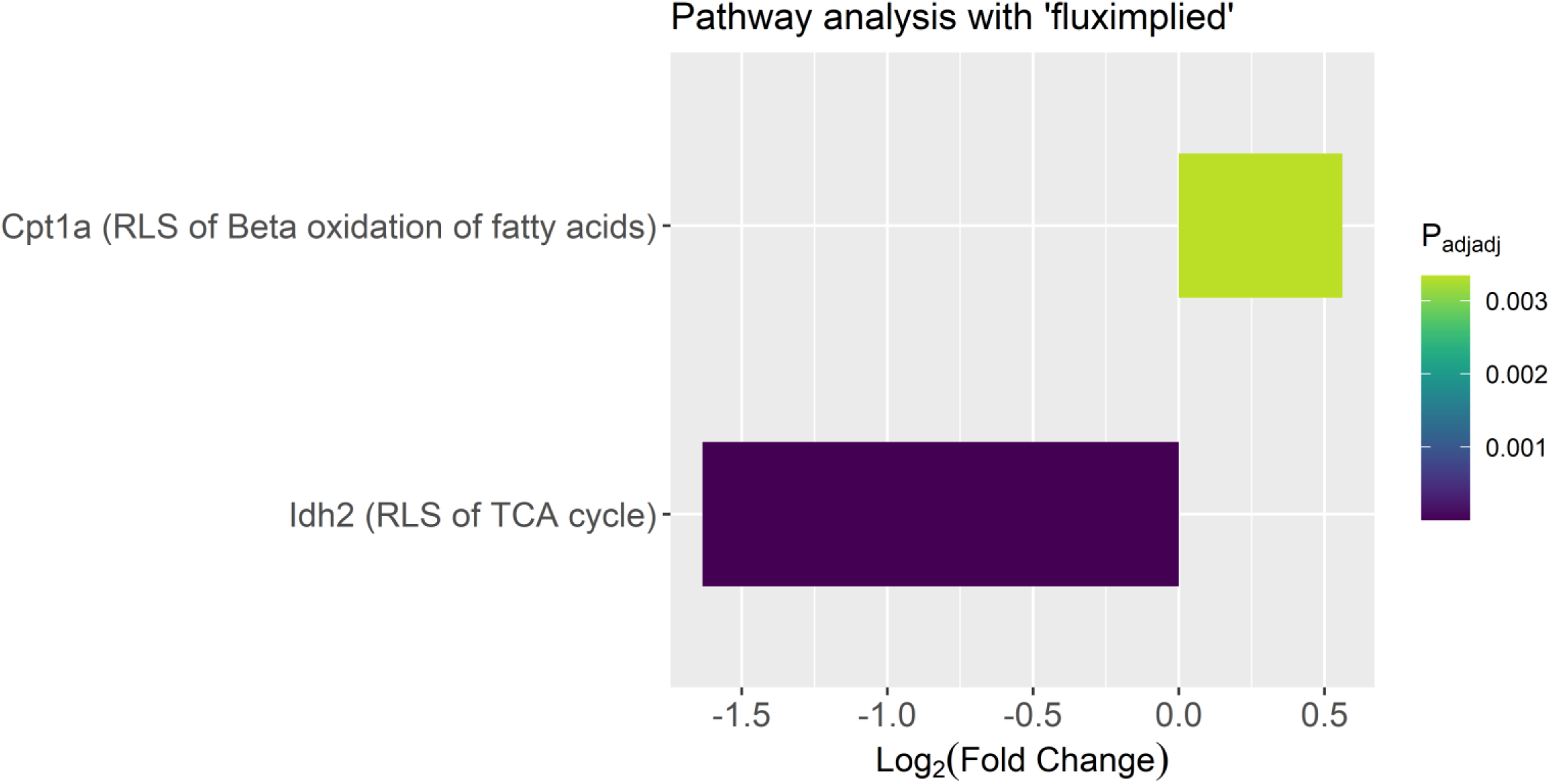
Example plot produced by flux implied. Fluximplied() will return a bar plot of each rate limiting step corresponding to its log_2_(fold change) colored by P_adjadj_.

If only a character vector of genes is supplied, then everything but the plot will be returned, but the significance table will include only your RLS genes supplied with their metabolic pathway and KEGG Pathway ID.

#### Example

Examples can be found in the vignette, which can be accessed at https://github.com/sportiellomike/fluximplied/blob/master/vignettes/fluximplied-vignette.Rmd or by going to the ‘help’ file in Rstudio and clicking ‘User guides, package vignettes and other documentation.’

Here are two examples for each kind of inputformat:

~~~
# data frame like input
inputdat<-exampleData
fluximplied(inputdat = inputdat,
species = ‘mmu’,
geneformat = ‘symbol’,
inputformat = ‘dataframe’,
padjcolname = ‘padj’,
pcutoff = .05
)
# vector like input
inputdat<-c(’Ifng’,‘Idh2′,‘Pfkl’)
fluximplied(inputdat = inputdat,
species = ‘mmu’,
geneformat = ‘symbol’,
inputformat = ‘vector’
)
~~~

### Web-based application

A web-based application is also provided for a responsive, interactive experience: https://sportiellomike.shinyapps.io/fluximplied/ (**Figure 3**).

**Figure 3:**
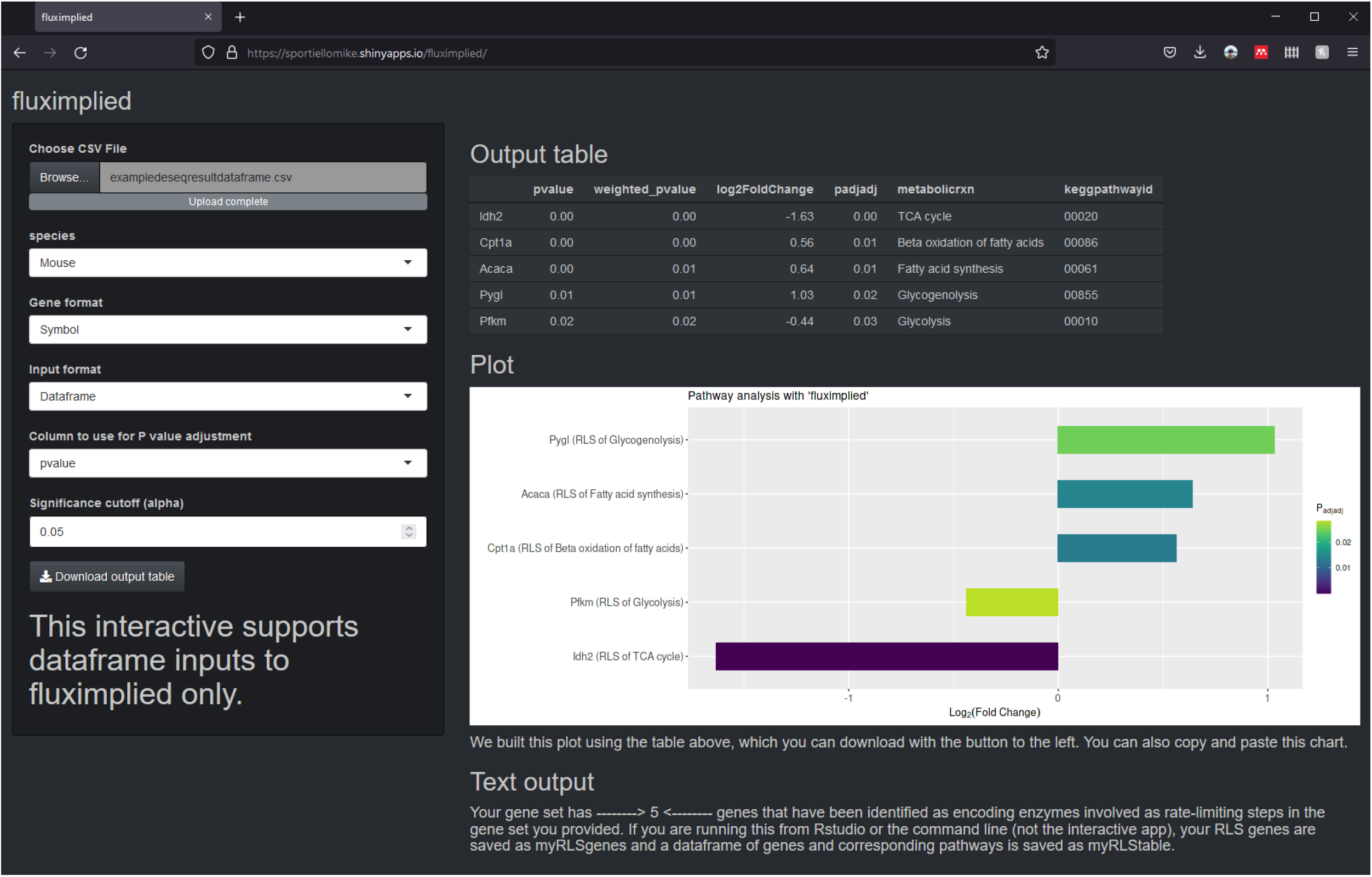
Interactive web application fluximplied. The web application is immediately responsive to changes in parameters to allow users to easy manipulate settings of fluximplied.

#### Inputs

Only CSVs, which will be treated like data frames after upload, are accepted (not character vectors of genes). This responsive GUI will update automatically after any parameter is changed, including species, gene format, which column of the uploaded CSV to treat as P_adj_, and the significance cutoff (only results with P_adjadj_ lower than this cutoff will be returned).

#### Outputs

The significance table, plot, and text to aid in interpretation are all updated automatically. The user may also copy and paste the plot, as well as download the significance table by clicking the download button.

## Results and Discussion

### Analysis of human tissues

In order to assess fluximplied’s utility we derived three bulk RNAseq datasets from the publicly available GTEx database. Specifically, we focused on three human tissue types with different metabolic capacities and requirements: subcutaneous adipose tissue, liver, and the putamen i.e. neuronal tissue. We performed differential expression analysis using DESeq2 for the three possible pairwise comparisons between the tissue datasets. DESeq2 output yields per-gene LFC and associated p values regarding differential expression as well as adjusted p values after correction for multiple comparisons and the use of independent hypothesis weighting^13,14^. Pathway analysis was performed on these datasets using EnrichR to query both the Reactome and KEGG databases^15^. In parallel, we used fluximplied to assess the differential expression of RLS transcripts between our tissue types as a form of validation.

EnrichR pathway analysis performed with both KEGG and Reactome yielded significant overlap with metabolism gene sets, particularly among liver transcripts relative to adipose and neuronal tissue. The fluximplied output was generally in concordance with typical enrichment analysis methods, but also identified physiologically valid differences in RLS transcript levels. For example, liver RNAseq datasets were enriched for *FPB1* and *CPS1* mRNA transcripts (the RLSs of gluconeogenesis and the urea cycle, respectively) relative to both adipose and neuronal tissue, reflecting the key roles of the liver in the regulation of blood glucose and the elimination of ammonia waste (**Figs. 4 and 5**) ^16^. Liver datasets were also enriched for *ACACA* but not *CPT1A* transcripts (the RLSs of fatty acid synthesis and β-oxidation, respectively) compared to adipose tissue. (**Figs. 4 and 5**) These findings are generally consistent with the liver’s relevance to lipid metabolism as a site for the synthesis, storage, and degradation of fatty acids according to the metabolic needs of the body ^16^. Adipose tissue is similarly capable of homeostatic metabolism of fatty acids, with synthesis generally occurring in the fed state and β-oxidation increasing with fasting or for the purpose of thermogenesis, according to the specific tissue type^17^.

**Figure 4:**
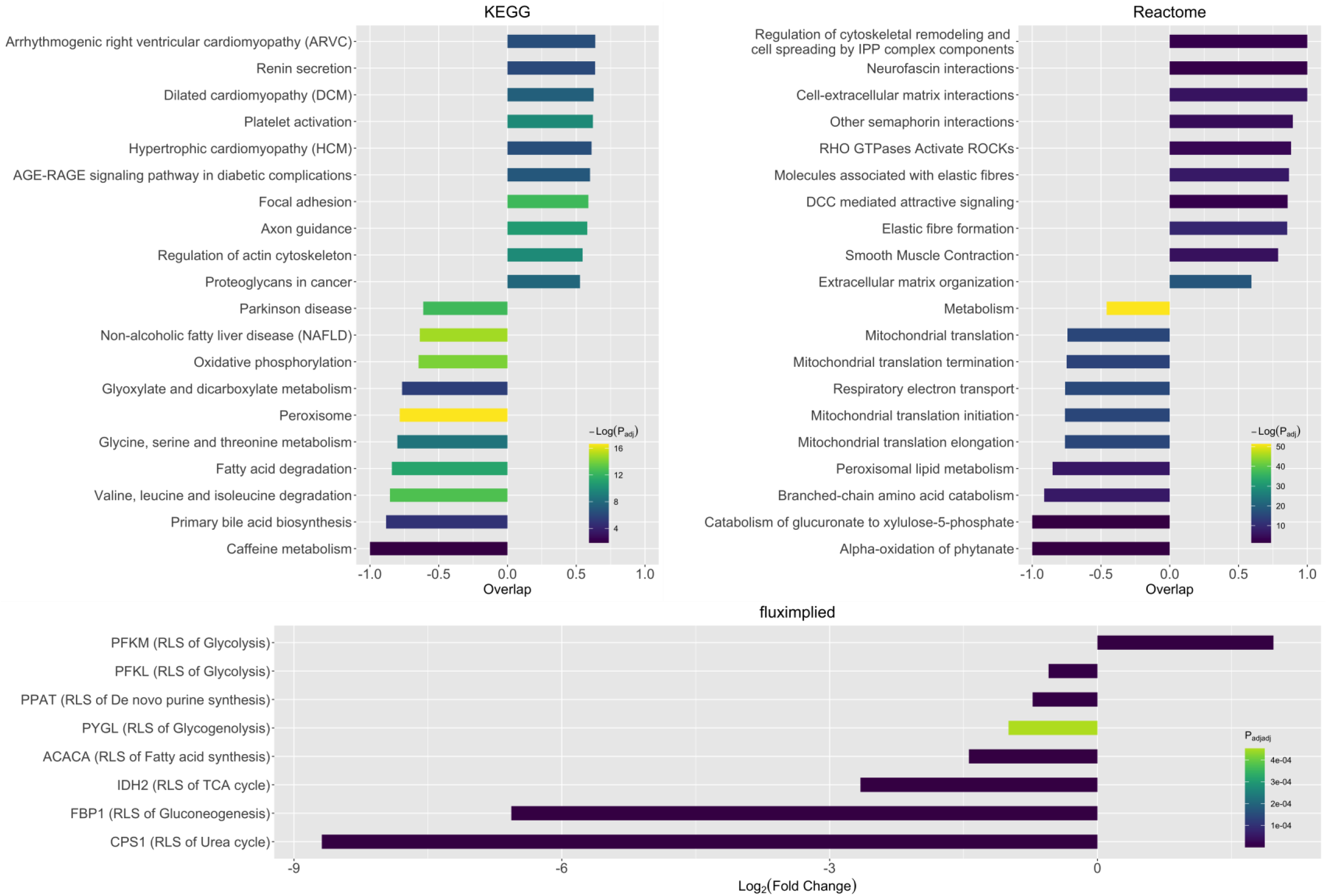
Adipose-liver differential expression analysis. Pathway Analyses for KEGG (A) and Reactome (B) databases. Overlap scores are calculated as the fractional number of genes present in a given gene set; negative overlap scores indicate downregulation for a given pairwise comparison. Gene sets were chosen for plotting by EnrichR’s combined score value: the top and bottom ten scoring sets for each comparison were ordered by overlap score. Fluximplied plot output (C). Hits to the fluximplied database are ordered by LFC value; negative values indicate downregulation for a given pairwise comparison.

**Figure 5:**
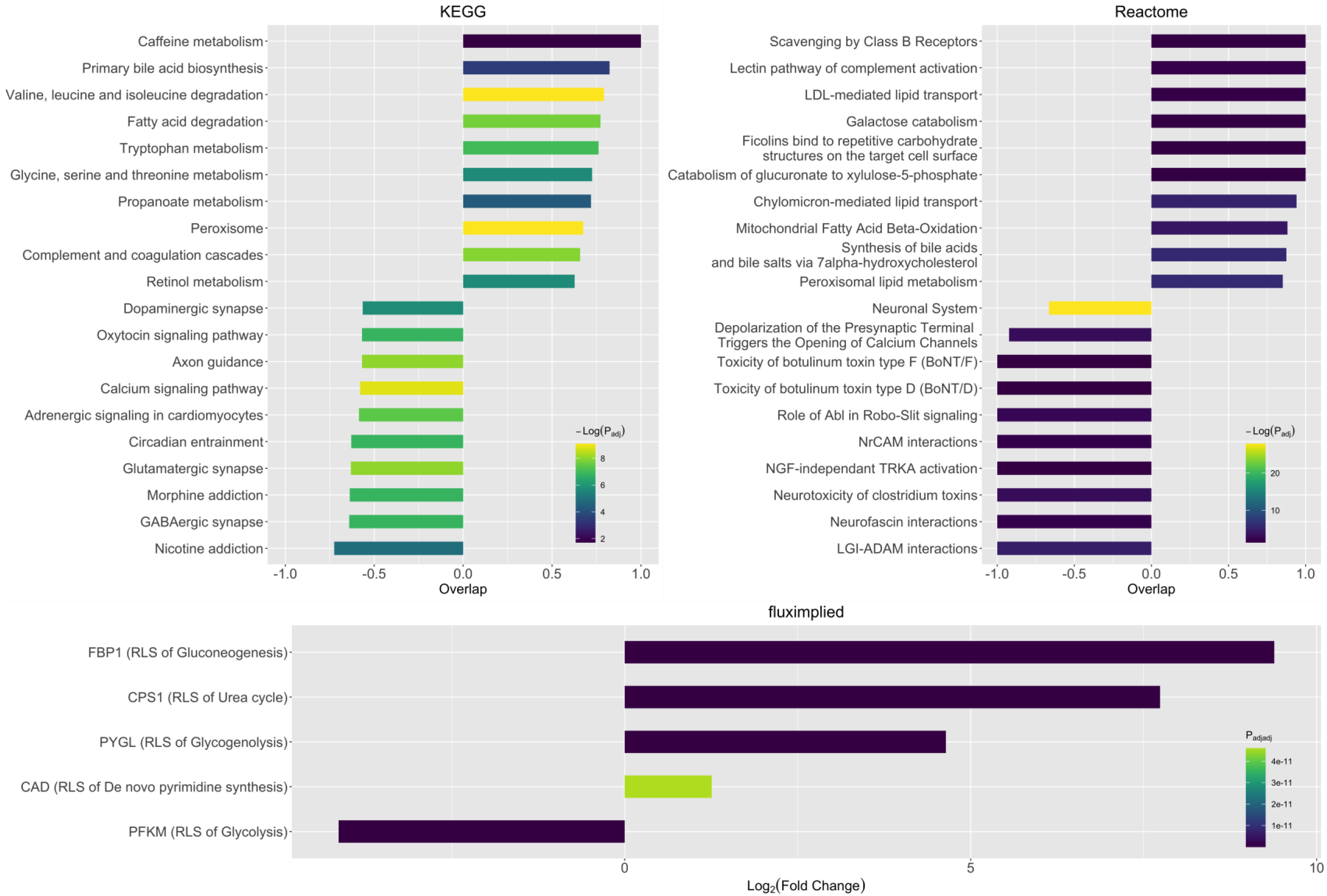
Liver-putamen differential expression analysis. Pathway Analyses for KEGG (A) and Reactome (B) databases. Overlap scores are calculated as the fractional number of genes present in a given gene set; negative overlap scores indicate downregulation for a given pairwise comparison. Gene sets were chosen for plotting by EnrichR’s combined score value: the top and bottom ten scoring sets for each comparison were ordered by overlap score. Fluximplied plot output (C). Hits to the fluximplied database are ordered by LFC value; negative values indicate downregulation for a given pairwise comparison.

Unlike liver and adipose tissue, neuronal tissue is known to be fundamentally dependent on glucose metabolism to meet its energetic needs ^18^. Among GTEx datasets, samples of the putamen showed the greatest degree of enrichment for *PFKM* transcripts (LFC = 2.138 and 4.135 relative to the adipose and liver tissue, respectively), implying a greater degree of glycolytic flux (**Figs. 5 and 6**). Recent evidence suggests that neuronal metabolism is more complex but our data demonstrate the fundamental reliance of central nervous tissue on glycolysis ^19^.

**Figure 6:**
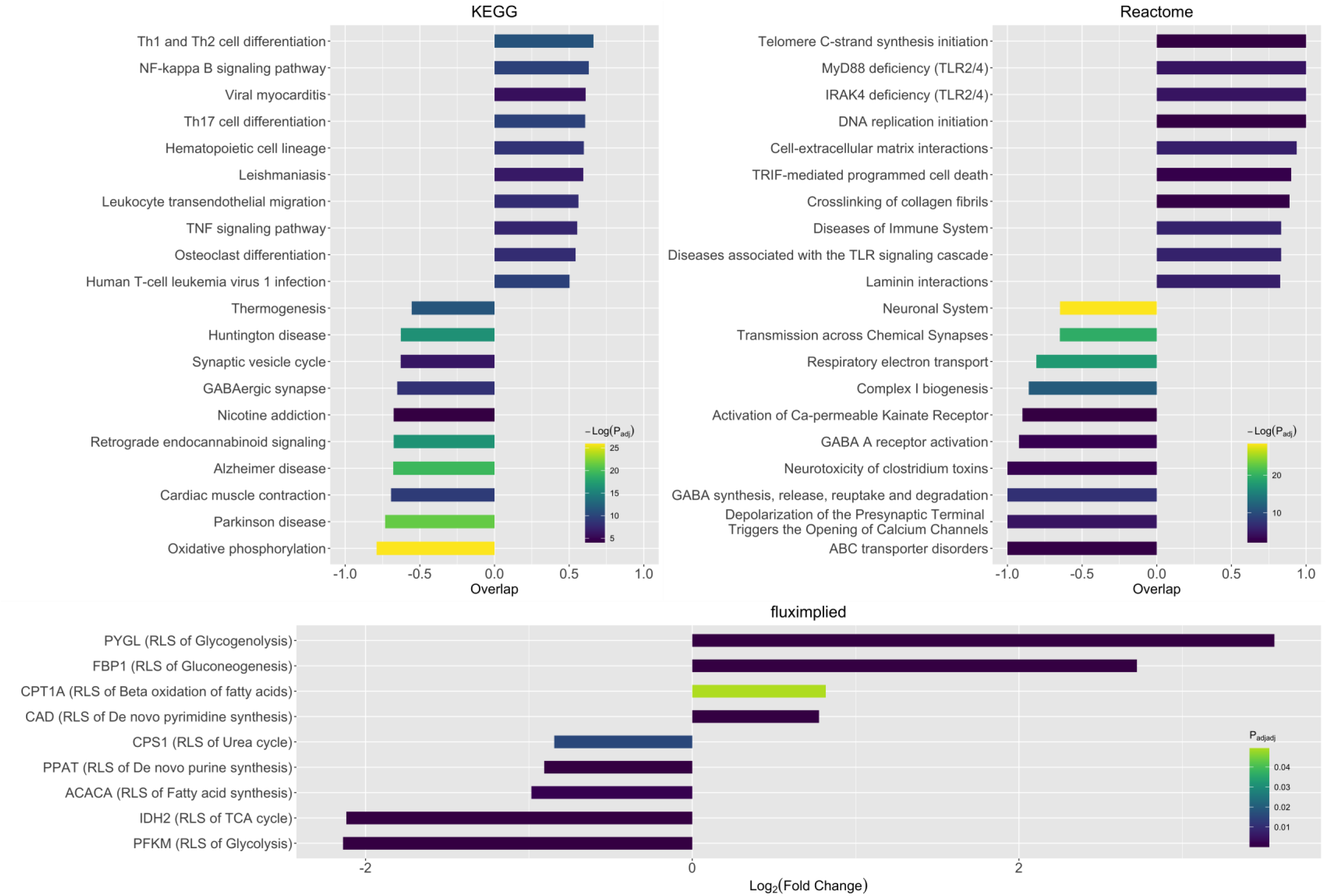
Adipose-putamen differential expression analysis. Pathway Analyses for KEGG (A) and Reactome (B) databases. Overlap scores are calculated as the fractional number of genes present in a given gene set; negative overlap scores indicate downregulation for a given pairwise comparison. Gene sets were chosen for plotting by EnrichR’s combined score value: the top and bottom ten scoring sets for each comparison were ordered by overlap score. Fluximplied plot output (C). Hits to the fluximplied database are ordered by LFC value; negative values indicate downregulation for a given pairwise comparison.

### Analysis of mouse lung CD8 T cell bulk RNAseq after influenza A virus infection

To validate fluximplied using data whose metabolic profiles are closer and therefore more difficult to interrogate, we used previously generated bulk RNAseq data from two memory CD8 T cell populations that come from the same mouse lungs and differed, at the time of sort, only in the expression of integrins CD49a and CD103 (GSE179653). In antigen-experienced cells, CD49a and CD103 are two markers of tissue resident memory while CD49a^−^CD103^−^ cells are generally considered circulating memory T cells. After preprocessing, these data were analyzed as described above for human tissue RNAseq datasets. Our results indicated that lung T_RM_ have a lipid centric metabolism as compared to circulating T cells, which had been demonstrated in skin. Synthesizing these findings, we set out to see if fluximplied could corroborate these findings.

Fluximplied reproduced what is broadly accepted about T_RM_: it identified β-oxidation of fatty acids as one of the main pathways whose RLS is upregulated (P_adjadj_<0.05)^20^ (**Figure 7**), which corroborates the findings of the metabolic model constructed in our recent publication (Sportiello *et alia*, submitted). Fluximplied was able to discover these pathways using the stricter LFC cutoffs of 0.5 and −0.5, which the typical enrichment analysis could not using the KEGG database, but could using the Reactome database. Not only was this technique more sensitive, fluximplied also generated a new hypothesis to be further investigated, as glycogenolysis was the pathway whose RLS was more upregulated than any other (LFC = 1.03). Though glycogenolysis has been reported in memory T cells before, it has never been investigated for functional importance in T_RM_.

**Figure 7:**
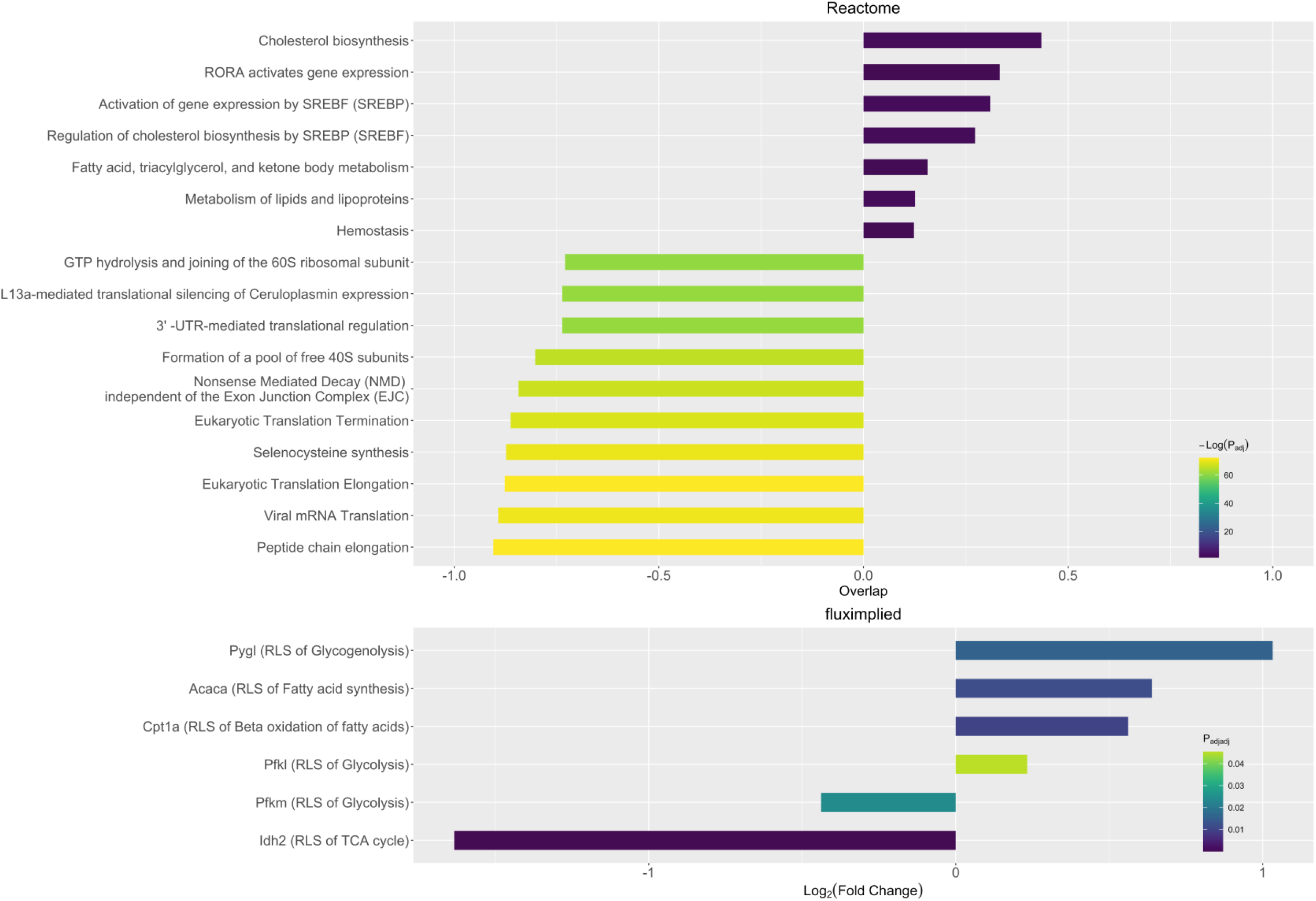
Mouse CD8^+^ tissue resident memory-circulating T cell differential expression analysis. Pathway Analyses for Reactome (A) database. Overlap scores are calculated as the fractional number of genes present in a given gene set; negative overlap scores indicate downregulation for a given pairwise comparison. Gene sets were chosen for plotting by EnrichR’s combined score value: the top and bottom ten scoring sets for each comparison were ordered by overlap score. Fluximplied plot output (B). Hits to the fluximplied database are ordered by LFC value; negative values indicate downregulation for a given pairwise comparison.

## Conclusions

In summary, we have built the hypothesis-generating tool fluximplied and demonstrated its concordance with other methods in human tissue. Next, we further demonstrated concordance and increased sensitivity using two memory CD8 T cell subsets using RNAseq datasets taken from biologically paired samples. Comparing two subsets of CD8 T cells from the same mouse lung demonstrate that this method is powerful. Finally, we demonstrated its ability to generate new hypotheses for data that contain less obvious differences in metabolic profiles.

There are immediate caveats for the interpretation of fluximplied’s output, most notably the presence of post-translational allosteric modification as a means of altering metabolic flux in many pathways. These regulatory processes would not be expected to alter transcriptional dynamics and therefore would not be integrated into fluximplied’s output; this is a problem with any use of transcriptomic data that could be ameliorated by use of proteomic data with fluximplied. Additionally, characterizations of reaction rate and flux are theoretical descriptions of complex biological processes and RLS designations may be reaction condition-dependent. In general, however, fluximplied can be seen as a tool for hypothesis generation and an extension of typical pathway analysis methodologies.

Currently, the RLSDatabase remains small, as one strength of the software is its power to find differences that are missed by traditional enrichment analyses and adding a plethora of pathways to it would lead to a decrease in power due to the multiple comparisons adjustment. Future directions include building out discrete tests for the user to select before running fluximplied, allowing the user greater control of pathways of interest without decreasing the program’s power. Expanded annotation of pathways that are not only RLSs, but also known regulatory enzymes due to post-translational modification would be of great benefit. Lastly, and maybe of most interest, investigating the interaction between metabolic pathways and immune pathways (termed “immunometabolic pathways”), would be of great use as well: many metabolites are important signaling molecules in immune processes^21^, and predicting their buildup due to a downregulation of an RLS downstream from their production would lead to a cell intrinsic immunomodulation, which may lead to further autocrine, paracrine, and endocrine signaling. This example regarding immunometabolism would hold true for other processes for which metabolites are known to participate, including DNA methylation^22^, memory formation^23^, and more.

## Supporting information

RLS Database

Up-reactome Mouse

Down-reactome mouse

Downregulated-Adipose vs Liver Kegg

Downregulated-Adipose vs Liver Reactome

Upregulated-Adipose vs Liver Kegg

Upregulated-Adipose vs Liver Reactome

Downregulated-Adipose vs Putamen Kegg

Downregulated-Adipose vs Putamen Reactome

Upregulated-Adipose vs Putamen Kegg

Upregulated-Adipose vs Putamen Reactome

Downregulated-Liver vs Putamen Kegg

Downregulated-Liver vs Putamen Reactome

Upregulated-Liver vs Putamen Kegg

Upregulated-Liver vs Putamen Reactome

## Availability and requirements

**Project name:** fluximplied

**Project home pages:**

- https://sportiellomike.shinyapps.io/fluximplied/
- https://github.com/sportiellomike/fluximplied

**Operating system(s):** Platform independent

**Programming language:** R

**Other requirements:** R 3.1 or higher

**License:** MIT License

**Any restrictions to use by non-academics:** N/A

## List of abbreviations

RLS: Rate Limiting Step
GSEA: Gene Set Enrichment Analysis
LFC: Log_2_(Fold Change)
KEGG: Kyoto Encyclopedia of Genes and Genomes
GUI: Graphical User Interface
P_adj_: adjusted P value after a differential expression analysis
P_adjadj_: fluximplied adjusts the adjusted P value to created the adjusted adjusted P value
CSV: comma separated value file, a standard format accessible to number of software including excel, LibreOffice Calc, and other text editors like Sublime Text, and Notepad.

## Declarations

- Ethics approval and consent to participate

- All data used are from human participants that consented to participate in the GTEx Portal or from publicly available mouse studies reviewed and approved by Institutional Animal Care and Use Committee of the University of Rochester.
- Consent for publication

- Not applicable
- Competing interests

- We report no competing interests.
- Funding

- Funding for this project came from NIAID P01-AI102851 which funded the experimental work; T32-HL066988 and T32-GM007356 training grants that fund MS, the first author of the manuscript.
- Authors’ contributions

- MS conceived fluximplied. MS and AG developed and validated the software. MS, AG, and RP wrote the manuscript. AKE, NGL, ECR, RP, and KLE provided critical advice and technical support for development of fluximplied. DJT oversaw the project, and secured funding for the project. All authors participated in critical editing of the manuscript.
- Acknowledgements

- The Genotype-Tissue Expression (GTEx) Project was supported by the Common of the Office of the Director of the National Institutes of Health, and by NCI, NHGRI, NHLBI, NIDA, NIMH, and NINDS.

## Notes

https://github.com/sportiellomike/fluximplied

### Competing Interest Statement

The authors have declared no competing interest.

